# Context-dependent transcriptional regulation of microglial proliferation

**DOI:** 10.1101/2021.03.29.436988

**Authors:** Sarah Belhocine, Andre Machado Xavier, Félix Distéfano-Gagné, Stéphanie Fiola, Serge Rivest, David Gosselin

## Abstract

Microglia proliferation occurs during brain development and brain lesions, but how this is coordinated at the transcriptional level is not well understood. Here, we investigated transcriptional mechanisms underlying proliferation of mouse microglia during postnatal development and in adults in models of induced microglial depletion-repopulation and brain demyelination. While each proliferative subset displayed globally a distinct signature of gene expression, they also co-expressed a subgroup of 1,370 genes at higher levels than quiescent microglia. Furthermore, expression of these may be coordinated by one of two modes of regulation. A first mode augments expression of genes already expressed in quiescent microglia and is subject to regulation by Klf/Sp, Nfy, and Ets transcription factors. Alternatively, a second mode enables de novo transcription of cell cycle genes and requires additional regulatory input from Lin54 and E2f factors. Overall, proliferating microglia integrate regulation of cell cycle gene expression with their broader, context-dependent, transcriptional landscape.

## Introduction

Microglia are the tissue resident macrophages of the central nervous system (CNS)^1^. These immune cells actively participate to CNS development and throughout life, promote its functions, protection against infections and ensure its repair following injuries. It is imperative that an optimal number of microglia be attained and maintained in any given context for microglial functions to be efficient. To this end, microglia can readily enter cell cycle and proliferate to increase its effective cell population in a variety of physiological settings. For example, work in mice has shown that during fetal and early post-natal development, microglia proliferate massively, mirroring the development and expansion of the brain^2,3^. In adults, the microglial population remains relatively stable, owing to an equilibrium between microglial cell death and cell proliferation^4,5^. Lastly, and perhaps most strikingly, microglia will promptly proliferate upon detection of pathogens, acute injuries and in chronic neurodegenerative disease^6–8^.

A cell’s ability to proliferate depends upon the transcriptional induction and repression of several genes at each of the different stages of the cell cycle process^2,9^. Triggering and coordinated these transcriptional changes requires regulatory input from numerous molecular effectors. With respect to microglia, evidence indicates that high stimulation of the colony-stimulating factor 1 receptor (Csf1r) microglia is likely to be an essential event responsible for initiating the cell cycle transcriptional program of these cells^10,11^. Indeed, as a tyrosine kinase receptor, Csf1r modulates many intracellular signaling pathways that possess potent transcriptional regulatory properties, including PI3K/Akt, SFK, MEK, JNK, ERK1/2^12^. Of note, these have previously been linked to cell cycle regulation in mononuclear phagocytes^12^. However, little is known about the identity of the transcription factors that provide regulatory input onto the microglial transcriptional machinery to coordinately effectively expression of cell cycles. While early evidence supported roles for transcription factors Runx1, Pu.1 and C/ebpβ^11,13^, the contribution of these as cell cycle regulators as opposed to factors that promote and maintain survival and general functionality of microglial has yet to be systematically established. Furthermore, and consistent with enhanced proliferative activities, subsets of microglia during brain development and brain lesions upregulate mRNAs coding for E2f family members, which are broad transcriptional regulators of cell cycle genes^2,9^. However, functional implication of E2f factors in microglia is still lacking. Lastly, studies in primary macrophages suggested that joint inhibition of expression of MafB and c-Maf factors elicits potent proliferative properties in these cells, suggesting that these factors may actively repress cell cycle transcriptional mechanisms^14^. That being said, absence of MafB in microglia does not appear to deregulate their proliferative capabilities during brain development^2^.

Efficient regulation of gene transcription also requires coordinated interactions between transcription factors and chromatin^15^. To large extent, interactions that provide functional input onto the transcriptional machinery take place at genomic regulatory elements, including promoters and promoter-distal elements (sometimes referred to as “genomic enhancers”). The former are the genomic sites where the transcriptional machinery required for effective gene transcription fully assembles. In contrast, promoter-distal elements provide the majority of a cell’s genomic binding sites for transcription factors acting in context-dependent fashion; these elements are thus important to specify space- and timing parameters of gene transcription^16^. Of interest, the systematic characterization of a cell’s repertoire of genomic regulatory elements is a sound strategy to identify transcription factors that potentially contribute to the transcriptional output of a cell in a given context. For example, promoter-distal elements that are preferably active in a given tissue-resident macrophage subset are enriched for binding sites recognized by transcription factors important to the functions of that macrophage population^17,18^. Thus, microglia display relative enrichment for Smad and Mef2 factors, large peritoneal macrophages are linked to Gata6 and Kupffer cells to Lxra. Notably, an increasing number of studies have validated functional roles for these factors or their regulators in coordinating the activity of their corresponding macrophage subset^19–21^.

As noted above, microglia proliferate in a variety of distinct physiological settings, each associated with its own molecular signature and constellation of cellular effectors. However, it is currently unknown whether similar or distinct programs of gene expression coordinate microglial proliferation across such varied settings. Indeed, the distinct signals associated with each setting may converge on a unique network of genomic regulatory elements, or alternatively, they may activate distinct networks each capable of mediating efficient cell cycling. Here, we examined this question using paradigms in epigenomics and, in parallel, also aimed to identify potential transcription factors that may contribute to the regulation of genes required for microglia to complete the cell cycle process efficiently.

## Results

### Ki67-RFP mice enable identification and isolation of microglia engaged in cell cycle

We used the Ki67^RFP^ mouse line to investigate transcriptional regulation of proliferative microglia. In this transgenic mouse, the gene coding for the Red Fluorescent Protein (RFP) reporter protein is fused in frame to the C-terminus end of the *Mki67* gene^22^, which is highly expressed in a do novo fashion in proliferating cells^23^. Thus, the RFP signal directly reflects abundance of the Ki67 protein. To validate efficacy of the RFP reporter in discriminating Ki67^+^ from Ki67^-^ microglia, we first analyzed with flow cytometry microglia isolated from various brain contexts previously characterized as been associated with microglial proliferation. These included postnatal day (PND) 8 neonates^24^ and adult mice (~PND80) sacrificed 3 days after undergoing 17 days of PLX5622 diet^25^. In this latter model, ingestion of rodent chow mixed with PLX5622, a potent Csf1r inhibitor, causes chronic depletion of more than 90% of microglia^25^. Upon return to regular chow diet (i.e., termination of PLX5622 administration), surviving microglia rapidly enter cell cycle, proliferate massively, and re-establish the microglial population to pre-PLX5622 diet level within one week^25^. Microglia from healthy, 60-75 days old adult mice were also included as negative controls, as most microglia in this condition are in a state of quiescence.

Microglia were defined as CD11b^+^CD45^+^CD44^Low^-single live cells (Fig. S1A). Overall, less than 2% of microglia from the healthy adult mouse brain emitted a RFP signal, which is coherent with a low proliferative rate in homeostatic condition (Fig. 1A)^4^. In contrast, both Ki67^-^ and Ki67^+^ microglial subsets populate the brains of PND8 neonates and those of mice 3 days after termination of the PLX5622 diet (Fig. 1A). Furthermore, staining of the Ki67 protein with anti-Ki67 antibody revealed that the Ki67^+^ antibody signal is highly concordant with the Ki67-RFP^+^ signal for both contexts of proliferation (Fig. S1B). As a very dim Ki67^+^ antibody signal was detected in microglia from adult healthy brains (Fig. S1C; *Adult-Basal* condition), these results strengthen our confidence that the presence of positive/high RFP signal in Ki67^RFP^ mice is indicative of high Ki67 protein expression.

**Figure 1.**
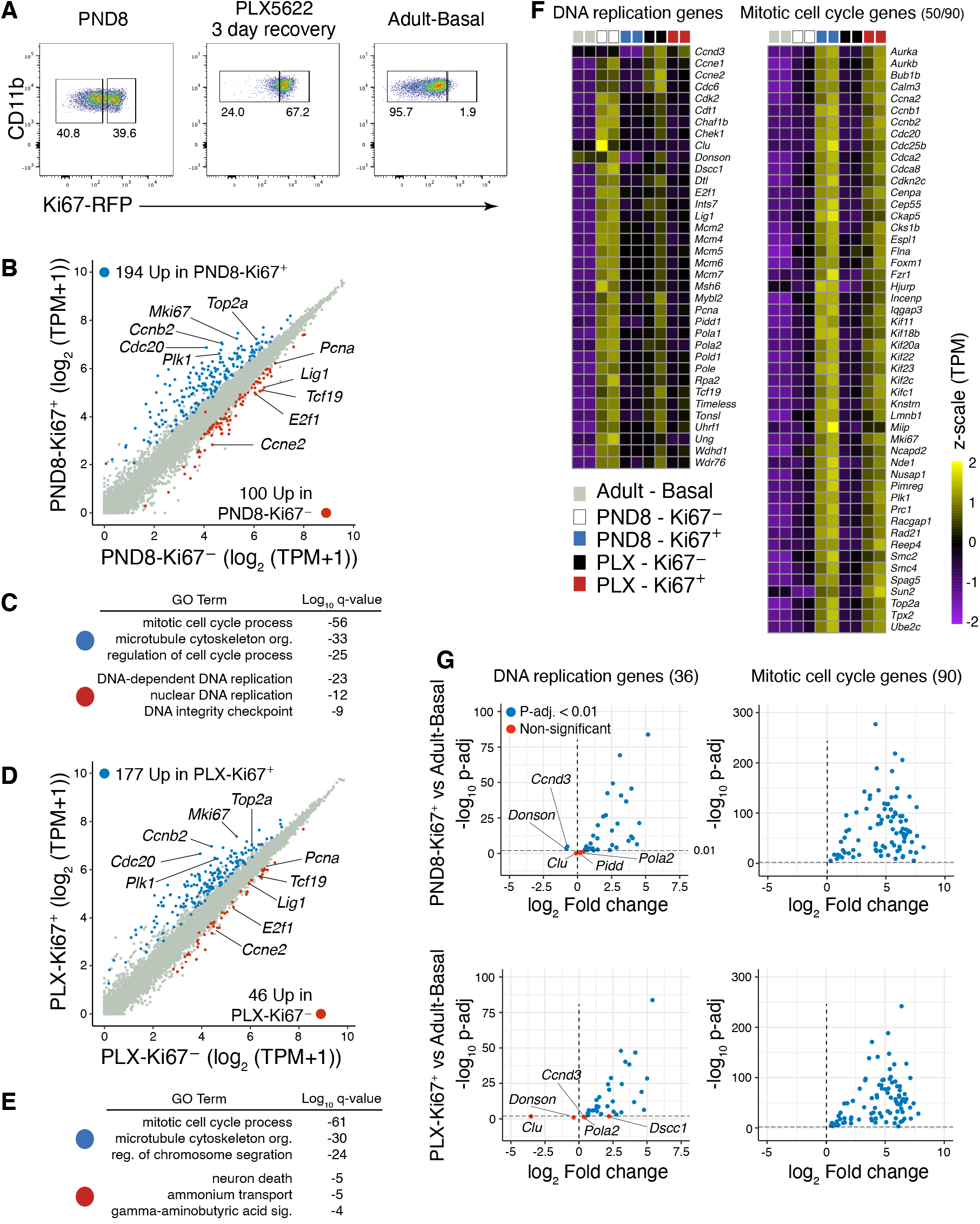
Ki67^+^ microglia express genes associated with DNA replication and mitosis. **(A)** Flow cytometry data depicting presence of Ki67^+^ microglia in the mouse brain during early postnatal brain development (PND8) and in adult three days after termination of administration of Csf1r inhibitor PLX5622. Ki67^+^ microglia are absent in the healthy, adult brain. Numbers represent percentage of microglia in corresponding gate above, relative to parent gate. **(B)** Scatterplot of RNA-seq data in Ki67^-^ and Ki67^+^ microglia isolated from the brain of postnatal day 8 (PND8) mouse neonates. Differentially expressed genes identified by DESeq2 (p-adj < 0.01) are colored coded. **(C)** Gene ontology of genes more highly expressed in PND8-Ki67^+^ vs PND8-Ki67^-^, and vice versa. **(D)** Scatterplot of RNA-seq data in Ki67^-^ and Ki67^+^ microglia isolated from the adult brain three days after termination of PLX5622 administration. Differentially expressed genes identified by DESeq2 (p-adj < 0.01) are colored coded. **(E)** Gene ontology of genes more highly expressed in PLX-Ki67^+^ vs PLX-Ki67^-^, and vice versa. **(F)** Heatmaps depicting z-normalized expression of gene, measured by RNA-seq, associated with GO categories *DNA-dependent DNA replication* (left) and *mitotic cell cycle process* (right). Each column represents independent biological replicates. **(G)** Volcano plots depicting fold change and p-adj. values of genes associated with GO categories *DNA-dependent DNA replication* (left column) and *mitotic cell cycle process* (right column), for the PND8-Ki67^+^ vs Adult-Basal (top row) and PLX-Ki67^+^ vs Adult-Basal (bottom row) comparisons.

We next characterized with RNA-seq the transcriptome of the Ki67^-^ and Ki67^+^ microglial subsets identified above. Relative to their Ki67^-^ counterparts, PND8-Ki67^+^ and PLX-Ki67^+^ microglia significantly expressed at higher levels 194 and 177 genes, respectively (false discovery rate (FDR) set at 1%; Fig. 1B and 1D, Table S1). These comprised numerous genes involved in the G2/M phase of the cell cycle, including *Mki67*, *Plk1*, *Ccnb2*. Alternatively, PND8-Ki67^-^ microglia from the neonatal brain expressed 100 genes more highly than their paired Ki67^+^ subset (FDR = 1%). Among these are genes required for DNA synthesis, including *Lig1*, *Pcna*, regulator of cyclin-dependent kinase *Ccne2* and cell cycle-associated transcription factor *E2f1*. However, while PLX-Ki67^-^ microglia also expressed many of these genes more highly than PLX-Ki67^+^ microglia, less genes met threshold for statistical significance, as only 46 genes were more abundant in the former compared to the latter (FDR = 1%).

The results above suggested that genes expressed at elevated levels in the Ki67^+^ vs Ki67^-^ subsets and vice versa may partitioned according to different phases of the cell cycle. Notably, gene ontology (GO) analyses confirmed that both Ki67^+^ subsets are enriched for genes linked to the mitotic cell cycle process and other processes that pertain to the G2/M stage of the cell cycle (Fig. 1C, E). In contrast, at PND8, a significant proportion of the genes more expressed in Ki67^-^ microglia relative to the Ki67^+^ subset include genes involved in DNA replication. However, transcripts more highly expressed in PLX-Ki67^-^ vs PLX-Ki67^+^ microglia did not display, collectively, strong linkage for a specific process of cell biology (Fig. 1E). This is possibly a consequence of the relatively low fraction of genes detected that are more highly expressed in the former subset compared to the latter. Nevertheless, collectively these results suggest that the Ki67^-^ and Ki67^+^ subsets are differentially enriched for microglia that are at different stages of the cell cycle process, with a predominance of G2/M microglia within the Ki67^+^ population and presence of G1/S microglia in the Ki67^-^ fraction. We note that the latter would also necessarily include G0 microglia as well.

Closer examination of the transcriptional signature of Ki67^+^ microglia revealed that although this subset may be dominated by cells at the G2/M stage, it also contains numerous mRNA transcript coding for proteins involved in DNA replication (Figure 1F). Indeed, of the 36 genes that make up the ensemble of genes linked to the DNA replication GO term as defined by PND8-Ki67^-^ microglia, 31 are also significantly more highly expressed in each of PND8-Ki67^+^ and PLX-Ki67^+^ microglia compared to Ki67^-^ microglia from healthy adult mice (Fig. 1G; Table S1). Whether this transcriptional signature arises from the proximity of the G1/S stages to the G2/M, or because the Ki67^+^ subsets include a noticeable number of microglia that maintain high levels of Ki67 protein as they rapidly re-enter cell cycle after G2/M exit, or a combination of both possibilities, is not discernable at this time. Finally, all 90 *mitotic* genes were significantly more expressed in both Ki67^+^ subsets (Fig. 1F, G). Overall, these results indicate that as a whole, the Ki67^+^ microglial population is likely to capture a wide spectrum of transcriptional events implicated in coordinating cell proliferation, including G1/S and G2/M-related events. Hence, analyses of Ki67^+^ microglia may represent a sound strategy to gain insights into the transcriptional mechanisms that coordinate microglial proliferation in the brain.

### Gene signatures of proliferative microglia are coupled to the brain physiological milieu

Given that microglia proliferate in different contexts, we next sought to determine whether microglial proliferation requires attainment of a unique transcriptional signature that is common to all context of proliferation, or whether a proliferation gene program integrates itself with other programs concomitantly active in microglia in a given context.

Pairwise comparison of Pearson correlation coefficients of the gene profiles of the different Ki67 subsets indicated that each Ki67^+^ subset is each more similar to its Ki67^-^ counterpart than to one another (Fig. 2A). This suggested that the surrounding brain microenvironment may have a widespread influence on the regulation of gene expression in proliferative microglia. To examine this in more details, we directly compared the transcriptomes of PND8-Ki67^+^ to that of PLX-Ki67^+^ microglia. At a false discovery rate of 1%, 860 genes are more highly expressed in PND8-Ki67^+^, including *Igf1*, *Gas7*, and *Fcrls* (Fig. 2B; Table S2). Alternatively, 1266 genes are elevated in the PLX-Ki67^+^ subset, and these comprised *Axl*, *Irf7*, *Sqle*, *Dhcr24*. Notably, GO analysis revealed that while upregulated genes in PND8-Ki67^+^ microglia are overall only weakly associated with terms related to *ribosome biogenesis* and *blood vessel morphogenesis*, genes more highly expressed in PLX-Ki67^+^ microglia are strongly linked to processes relevant to innate immune responses (Fig. 2C). Thus, while both PND8-Ki67^+^ and PLX-Ki67^+^ microglia are clearly in a state of cell proliferation, it appears that this state is also compatible with additional, distinct cellular states and functions.

**Figure 2.**
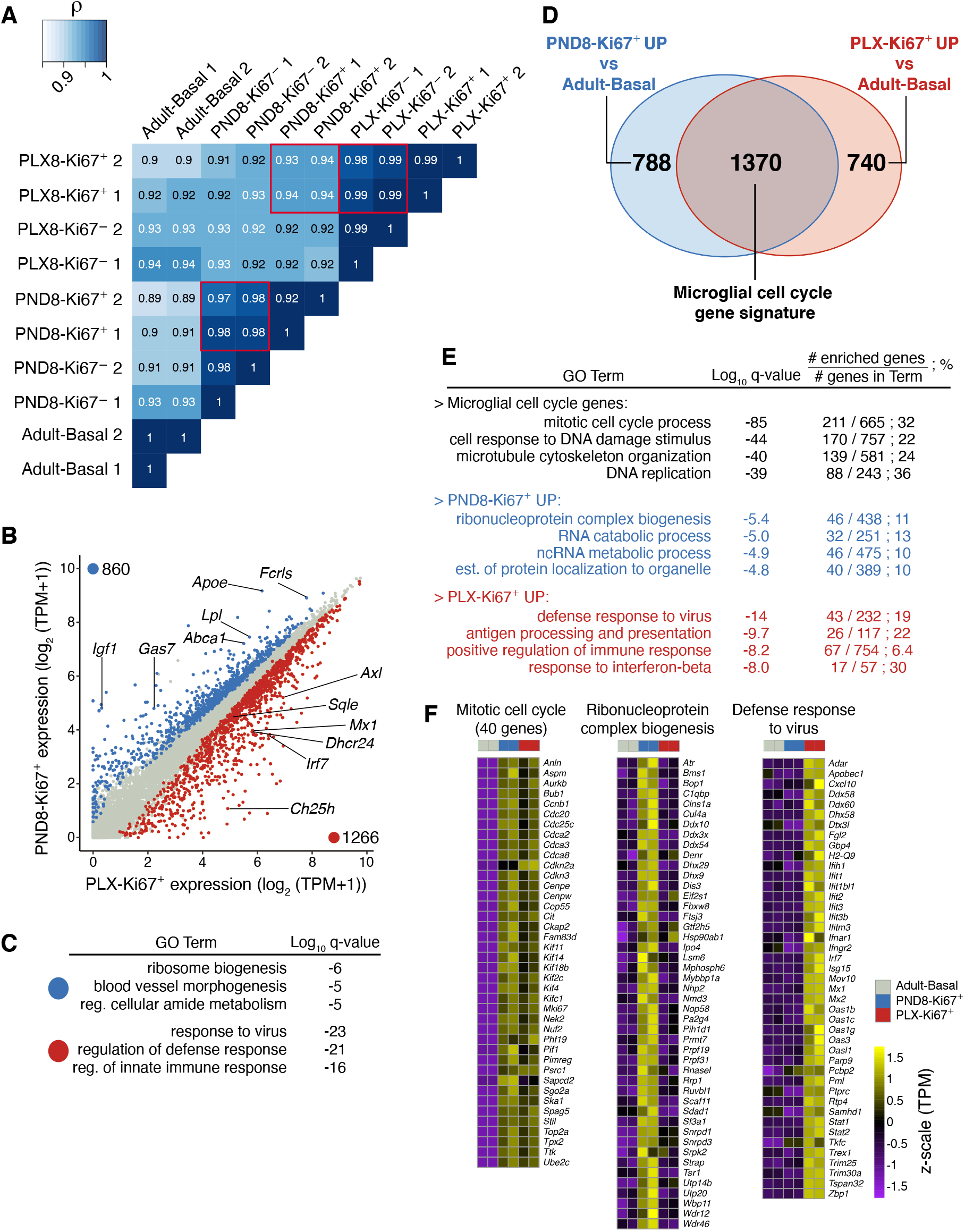
Transcription in proliferative microglia is coupled to the brain physiological state. **(A)** Pearson correlation data matrix, comparing Adult-Basal and Ki67^-^ and Ki67^+^ microglia from PND8 neonates and adults three days after termination of PLX5622 administration. **(B)** Scatterplot of RNA-seq data in PND8-Ki67^+^ and PLX-Ki67^+^ microglia. Differentially expressed genes identified by DESeq2 (p-adj < 0.01) are colored coded. **(C)** Gene ontology of genes more highly expressed in PND8-Ki67^+^ vs PLX-Ki67^+^, and vice versa. **(D)** Venn diagram representing genes significantly upregulated in PND8-Ki67^+^ vs Adult-Basal, and PLX-Ki67^+^ vs Adult-Basal (DESeq2 analysis; p-adj < 0.01). **(E)** Gene ontology of genes significantly induced in both PND8-Ki67^+^ and PLX-Ki67^+^ vs Adult-Basal (top), PND8-Ki67^+^ vs Adult-Basal only (middle) and PLX-Ki67^+^ vs Adult-Basal only (bottom). **(F)** Heatmaps depicting z-normalized expression values of gene, measured by RNA-seq, associated with GO categories *mitotic cell cycle process* (left), *ribonucleoprotein complex biogenesis* (middle) and *defense response to virus* (right). Each column represents independent biological replicates.

Though the transcriptomes of the PND8-Ki67^+^ and PLX-Ki67^+^ subsets do not mirror each other perfectly, it is possible that among all the transcriptional programs active in these cells a program necessary for *cell proliferation* is predominant. To verify this, we first recovered the genes that are significantly induced in each of the Ki67^+^ microglia subsets relative to G0/quiescent Ki67^-^ microglia isolated from the healthy adult brain (FDR = 1%). As this latter population is unlikely to include a sizeable fraction of proliferative microglia, we reasoned that they would provide a robust baseline to define context-dependent gene programs. Overall, 2,158 and 2,110 genes were upregulated in PND8-Ki67^+^ and PLX-Ki67^+^ microglia, respectively (Fig. 2D). Notably, a majority (1370, > 60%) of genes was common to both Ki67^+^ subsets’ signatures (Table S2). These *Common-Up* genes included a large ensemble of genes that are strongly associated with regulatory mechanisms of cell cycle, including processes of mitotic cell cycle and DNA replication (Fig. 2E, F). Alternatively, PND8-Ki67^+^-biased *Up* genes included genes linked to RNA regulation, while PLX-Ki67^+^-biased *Up* genes strongly associated with immune responses, which is consistent with results from the PND8-Ki67^+^ vs PLX-Ki67^+^ comparison.

Overall, these results indicate that microglia that proliferate in different physiological contexts can do so without reaching an identical transcriptional profile. That being said, these cells appear to devote substantial resources towards the transcription and preservation of gene transcripts involved in cell cycling. Therefore, proliferating microglia integrate a cell cycling gene program with multiple other programs, many of which are intrinsically linked to the brain microenvironment.

### Distinct combinations of transcription factors regulate different programs of gene expression associated with microglial proliferation

The *Common-Up* gene signature of microglial cell cycle identified above likely includes genes that are critical to ensure efficacious microglial proliferation in most, if not all, situations where microglial proliferation occurs. While examining the genes that make-up this *microglial proliferation* gene signature, we noticed that some genes are not expressed in Ki67^-^ Adult-Basal microglia but are robustly induced in the Ki67^+^ subsets (Fig. 3A). In contrast, other genes are well expressed in Ki67^-^ quiescent adult microglia under healthy condition but are induced at lower magnitude in Ki67^+^ microglia (Fig. 3A). Performing k-mean analysis based on these variables (i.e., expression levels in Ki67^-^ Adult-Basal vs fold-induction in Ki67^+^ states relative to Ki67^-^ Adult-Basal), partitioned genes of the *proliferation* signature into two distinct clusters. On the one hand, Cluster 1 (C1) comprised 881 genes, including *Dnmt1*, *Pcna*, *Cdk4*, and *Eef2*. Many of these predominately associated with cellular processes of translation, purine metabolism and mitotic cell cycling (Fig. 3B; Table S3). On the other hand, a large number of the 489 genes included in the second cluster (C2) were highly significantly linked to the mitotic cell cycle and DNA replication processes (Fig. 3B). Examples of C2 genes included *Mki67*, transcription factors *E2f1* and *Tcf19* and anaphase promoting complex member *Cdc20* (Fig. 3A). Of interest, although C2 genes are not or only very lowly expressed in Ki67^-^ quiescent microglia, the high abundance of the H3K4me3 histone mark encompassing their TSS in quiescent microglia suggests that the promoters of these genes are poised for transcriptional activation (Fig. 3C-E)^26^. Overall, these data indicate that multiple modes of transcriptional regulation coordinate gene expression in proliferating microglia. Thus, while a first mode (i.e., C1) augments the expression of genes linked to translation and metabolism that are already expressed in quiescent microglia, a second mode (i.e., C2) enables the de novo transcription of a large fraction of genes required for DNA synthesis and mitosis.

**Figure 3.**
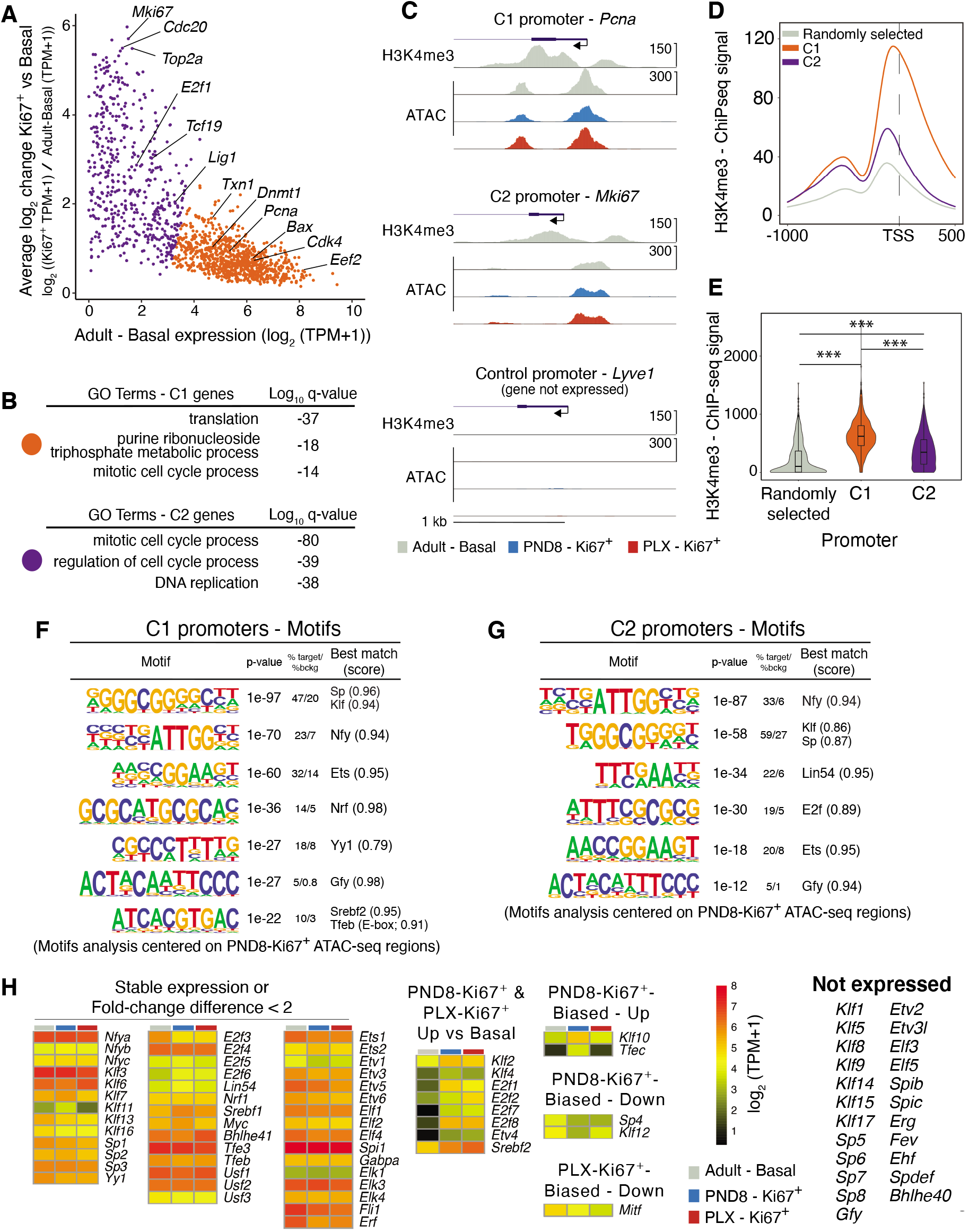
Distinct modes of transcriptional regulation coordinate gene expression in proliferative microglia. **(A)** Scatterplot depicting relationship between expression levels of microglia cell cycle genes in Adult-Basal condition vs average induction fold in PND8-Ki67^+^ and PLX-Ki67^+^. K-mean analysis was used to unbiasedly assign genes to Cluster 1 (C1, in orange) or Cluster 2 (C2, in purple). **(B)** Gene ontology of C1 and C2 genes (Metascape). **(C)** UCSC browser view of H3K4me3 and ATAC-seq data for representative C1 (*Pcna*), C2 (*Mki67*), and control (*Lyve1*) gene promoter loci. **(D)** Average H3K4me3 abundance over C1, C2 and control gene promoters, encompassing the −1000 to 500 bp region relative to transcriptional start sites. **(E)** Violin plots depicting quantification of the H3K4me3 reads over C1, C2, and control groups of gene promoters. Kruskal-Wallis test followed by Dunn’ test was performed to assess statistical significance of differences between the groups’ distribution of values (p<0.001(***)). **(F)** Tables summarizing results from de novo motifs analyses performed on C1 promoter regions. **(G)** Tables summarizing results from de novo motifs analyses performed on C2 promoter regions. **(H)** Heatmaps depicting gene expression values measured by RNA-seq of genes coding for transcription factors implicated by de novo motif analyses in (F) and (G) above.

We next investigated whether different transcriptional regulators may be implicated in controlling these transcriptional modalities. For this, we performed de novo DNA motifs analyses targeting promoters of C1 and C2 genes to identify candidate transcription factors that may drive transcriptional activity at these promoters. Specifically, regions included in these analyses consisted of areas of accessible chromatin, as defined by assay for transposase accessible chromatin sequencing (ATAC-seq), that are included within the −1000 to +500 bp regions encompassing the transcriptional start site (TSS) of each gene of each cluster. Of interest, analyses performed on each cluster returned both *cluster-common* and *cluster-specific* enrichments of motifs (Fig. 3F, G; Fig. S2). Thus, both C1 and C2 gene promoter regions displayed significant enrichment for motifs linked to the Klf/Sp, Nfy and Ets families of transcription factors. However, several C1 promoter elements appear to also be regulated by Yy1, and possibly members of the Ebox factor Srebf and Atf families, among others. In contrast, regulation of C2, but not C1, promoters possibly require contribution from Lin54 and E2f family members. Therefore, qualitatively distinct combinatory activity of transcription factors may promote gene transcription of C1 and C2 genes. As such, these data provide additional evidence that distinct modes of transcription may coordinate gene expression required for microglial cell cycling.

To gain a more precise understanding of which transcription factors may contribute to the regulation of the C1 and C2 gene promoters, we next assessed the mRNA expression levels of the factors implicated by the DNA motif analyses. Overall, 80 different transcription factors were recovered, 56 of which were reliably expressed in at least one dataset (i.e., TPM values of 16 or more in Ki67^-^-Adult-Basal or one of the Ki67^+^ subsets; Fig. 3H). Overall, the majority of these are either stably expressed across all microglial phenotypes or are only subtlety modulated in a context-dependent manner (i.e., less than two-fold change). However, among the 8 factors induced in both Ki67^+^ subsets compared to Ki67^-^ Adult-Basal microglia (two-fold increase, FDR = 5%), 4 are strongly upregulated by more than 8-fold. These include *E2f1* and Ets factor *Etv4*, which are part of the C2 gene cluster defined above. Thus, given the absence of, or the very low expression of these factors in basal condition, their induction might represent a key regulatory checkpoint for the engagement and/or efficient completion of microglial proliferation.

### Gain of activity of microglial promoter-distal genomic regulatory elements in Ki67^+^ proliferative microglia

We next analyzed the repertoire of promoter-distal regulatory elements of microglia to gain a more comprehensive understanding of the transcriptional mechanisms associated with microglial proliferation. For this, we interrogated with chromatin immunoprecipitation followed by massively parallel sequencing (ChIP-seq) the presence of the H3K27 acetylation mark (H3K27ac) on a genome-wide level, which enables assessment of relative activity at genomic regulatory elements^27^. We first compared profiles generated from PND8-Ki67^-^ to PND8-Ki67^+^ to validate that our ChIP-seq strategy can detect histone acetylation features associated with cell cycling. Overall, both profiles were similar to one another, with only 1% of the regions being differentially acetylated at H3K27 (350/32922; FDR 1%; Fig. S3A). Nevertheless, PND8-Ki67^+^ displayed significantly higher levels of H3K27ac at 124 promoter regions and these were enriched for genes implicated in the different stages of the cell cycle process, including G1/S and G2/M (Fig. S3B-S3E).

Based on these observations, we further analyzed Ki67^+^ microglia to identify possible promoter-distal genomic regions whose activity might help coordinate gene expression in proliferative microglia. For this, we compared the profiles of promoter-distal H3K27ac of the PND8-Ki67^+^ and PLX-Ki67^+^ subsets to that of Adult-Basal microglia. At a FDR of 1%, 1343 and 2576 regions were relatively more abundant for H3K27ac in Ki67^+^ microglia from PND8 neonates and the PLX model, respectively, out of total of 24,007 and 24,429 regions (Fig. 4A, B). In contrast, 3264 and 3457 regions had relatively lower levels of this mark compared to Adult-Basal microglia, again respectively. Overlapping regions recovered from PND8 and PLX for each of the differential trajectories yielded 536 shared regions that are *Common-Up*, and 1712 shared regions that are *Common-Down* (Fig. 4C-F). Thus, these results suggest that transcriptional events associated with microglial proliferation modulates H3K27ac deposition at over ~2,200 promoter-distal sites. Further comparison of H3K27ac levels at these sites between PND8-Ki67^+^ and PLX-Ki67^+^ revealed Pearson correlation coefficients of 0.78 for *Common-Up* and of 0.92 for *Common-Down* elements (Fig. 4G, H). The relatively lower coefficient for *Common-Up* elements compared to *Common-Down* may suggest that these sites are particularly subject to important context-dependent effects on a quantitative dimension. We note that this would be coherent with the key role of enhancers in regulating space and time parameters of gene expression^15^.

**Figure 4.**
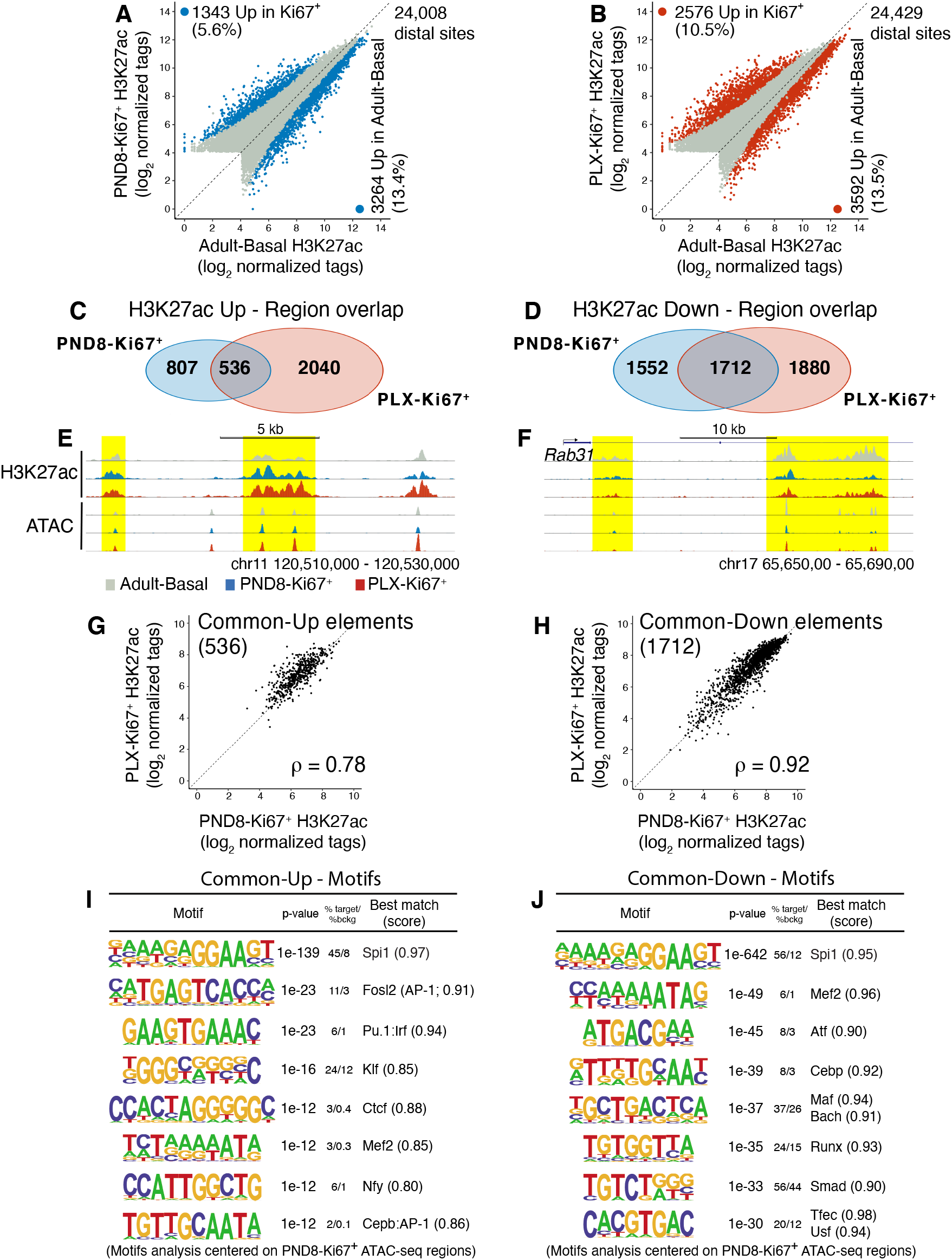
Differential H3K27ac abundance at promoter-distal genomic regulatory element associated with microglial proliferation. **(A)** Scatterplot representing H3K27ac abundance at promoter-distal genomic regulatory elements, comparing PND8-Ki67^+^ to Adult-Basal condition. Regions that display significantly higher H3K27ac signal are color coded (DESeq2 analysis, p-adj < 0.01). **(B)** Scatterplot representing H3K27ac abundance at promoter-distal genomic regulatory elements, comparing PLX-Ki67^+^ to Adult-Basal condition. Regions that display significantly higher H3K27ac signal are color coded (DESeq2 analysis, p-adj < 0.01). **(C)** Venn diagram representing promoter-distal regions with significantly higher levels of H3K27ac in PND8-Ki67^+^ vs Adult-Basal, and PLX-Ki67^+^ vs Adult-Basal (DESeq2 analysis; p-adj < 0.01). **(D)** Venn diagram representing promoter-distal regions with significantly lower levels of H3K27ac in PND8-Ki67^+^ vs Adult-Basal, and PLX-Ki67^+^ vs Adult-Basal (DESeq2 analysis; p-adj < 0.01). **(E)** UCSC browser view of H3K27ac ChIP-seq and ATAC-seq data for representative regions that gain H3K27ac in Ki67^+^ microglia vs Adult-Basal. **(F)** UCSC browser view of H3K27ac ChIP-seq and ATAC-seq data for representative regions that lose H3K27ac in Ki67^+^ microglia vs Adult-Basal. **(G)** Scatter plot depicting H3K27ac ChIP-seq signal for promoter-distal regions that gain acetylation at H3K27 in both of PND8-Ki67^+^ and PLX-Ki67^+^ vs Adult-Basal microglia. **(H)** Scatterplot depicting H3K27ac ChIP-seq signal for promoter-distal regions that lose acetylation at H3K27 in both of PND8-Ki67^+^ and PLX-Ki67^+^ vs Adult-Basal microglia. **(I)** De novo motifs enriched in open chromatin area located within promoter-distal regions that gain H3K27ac in Ki67^+^ microglia vs Adult-Basal. **(J)** De novo motifs enriched in open chromatin area located within promoter-distal regions that lose H3K27ac in Ki67^+^ microglia vs Adult-Basal.

We next performed DNA motifs enrichment analyses on *Common-Up* and *Common-Down* promoter-distal regulatory elements to identify candidate factors that may regulate gene expression by interacting with these sites to enable microglial cell cycling. For this, we again focused our analyses on areas of accessible chromatin defined by ATAC-seq data from PND8-Ki67^+^ that are encompassed within the larger regulatory elements of interest identified by presence of H3K27ac. These analyses revealed that for regions that gain H3K27ac, a few factors besides Ets family member Pu.1 and Klf family members appear to provide extensive regulatory input (Fig. 4I). Similar observations were made using ATAC-seq from PLX-Ki67^+^ microglia (Fig. S4A). Furthermore, elements that gain H3K27ac preferably in PLX-Ki67^+^ microglia displayed strong enrichment for motifs linked to AP-1 and Pu.1-Irf8 transcriptional regulators. This latter observation is highly consistent with the *response to virus* gene signature of this subset noted above and pro-inflammatory signaling pathways activated in this subset previously reported^28^ (Fig. S4B). In contrast, regions that are lose acetylation at H3K27 in Ki67^+^ microglia are enriched for motifs linked to Smad factors, among others (Fig. 4J). In particular, the Smad motif is also enriched at promoter-distal regions that lose H3K27ac in PND8-Ki67^+^ microglia (Fig. S4C). This observation suggests that microglial proliferation may require a decrease in Transforming growth factor beta (Tgf-β) regulatory input, as Smads are transcriptional effectors of this signaling system. This would be concordant with observations previously reported of enhanced microglial proliferation in absence of Tgf-β signaling in the brain^29^.

Together, these results indicate that different signaling pathways are enabled in parallel in proliferative microglia from different settings. This is coherent with the different gene signatures detected in these different proliferative microglia subsets. Moreover, there appears to be a relatively modest contribution for a gain of activity of a core set of promoter-distal elements in promoting transcription of gene transcription required for proliferation. In contrast, these results highlight that proper calibration of proliferation-inhibitory signals, such as Tgf-β/Smad activity, may be critically involved in coordinating efficient microglial proliferation.

### Microglia proliferating during brain demyelination display partial conservation of activity at genomic regulatory elements with PND8-Ki67^+^ and PLX-Ki67^+^ microglia

Microgliosis, which is characterized by extensive proliferation and accumulation of inflammatory microglia in the CNS, is an inherent feature of numerous neurodegenerative disorders and brain injuries. Such physiological contexts are not recapitulated in the postnatal brain or in the PLX5622 model of microglial depletion-repopulation. Thus, we next investigated the extent to which microglial proliferation associated with microgliosis proceeds through similar transcriptional mechanisms as those characterized above from Ki67^+^ microglia sampled from non-lesion contexts. To sample microgliosis-associated microglia, we isolated Ki67^-^ and Ki67^+^ microglia isolated from mice administrated cuprizone (CPZ) neurotoxin for 4 weeks through their diet. In this model, chronic ingestion of cuprizone induces heavy demyelination throughout the brain, which in turn is accompanied by substantial microgliosis^30^. Indeed, as shown in Fig. 5A, a clear subset of CD11b^+^CD45^+^CD44^Low^ microglia express high levels of Ki67 protein, as detected with anti-Ki67 antibody. Unexpectedly however, we were unable to detect RFP signal in Ki67^RFP^ mice fed cuprizone. Furthermore, as the cell permeabilization required for anti-Ki67 antibody labeling in combination with fluorescence-activated cell sorting dramatically damages RNA transcripts, we were not able to generate high quality RNA-seq gene profiles for CPZ microglial subsets. However, the anti-Ki67 labeling procedure is compatible with H3K27ac ChIP-seq (Fig. 5B).

**Figure 5.**
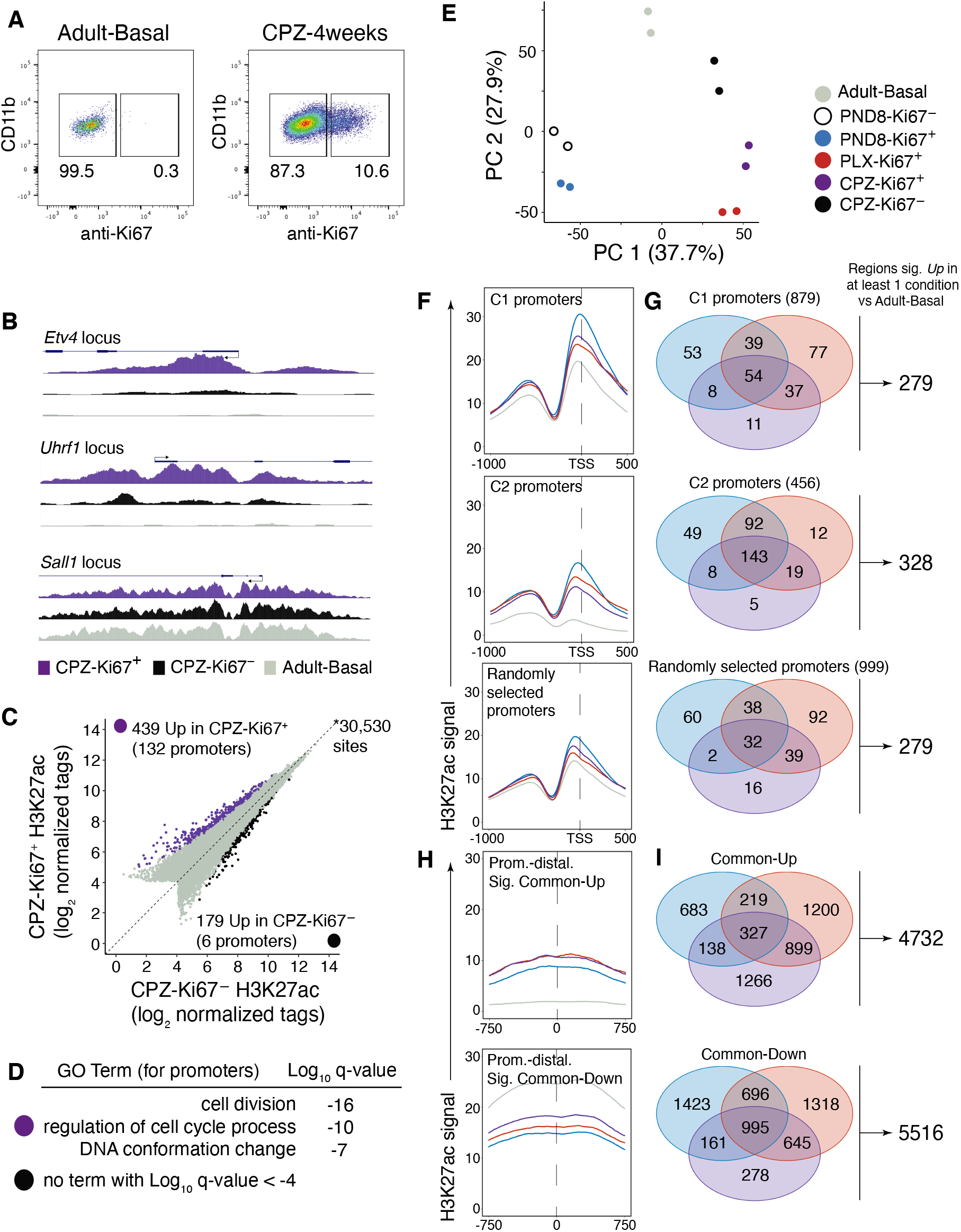
H3K27ac ChIP-seq analyses of proliferating microglia in the demyelinating brain. **(A)** Gating strategy for fluorescence-activated cell sorting-mediated isolation of Ki67^-^ and Ki67^+^ microglia from the brain of mice fed cuprizone (CPZ) for 4 weeks. Numbers represent percentage of microglia in corresponding gate above, relative to parent gate. **(B)** UCSC browser view of H3K27ac ChIP-seq and genes coding for cell cycle regulators (*Etv4*, *Uhrf1*) and control gene (*Sall1*) locus. **(C)** Scatterplot depicting H3K27ac abundance at genomic regulatory elements, comparing CPZ-Ki67^+^ to CPZ-Ki67^-^ microglia. Regions that display significantly higher H3K27ac signal are color coded (DESeq2 analysis, p-adj < 0.01). **(D)** Gene ontology analysis for promoter regions that display differential acetylation at H3K27 in CPZ-Ki67^+^ vs CPZ-Ki67^-^ microglia, and vice versa. **(E)** Principal component analyses of H3K27ac ChIP-seq data from Ki67^+^ and Ki67^-^ microglia isolated from various brain conditions. **(F)** Average H3K27ac abundance over the −1000 to 500 bp region surrounding the transcriptional start sites of C1, C2, and control gene promoters for different Ki67^+^ microglial subsets and Ki67^-^ microglia from healthy adult mice. **(G)** Venn diagram depicting overlap of among C1, C2 and control promoters that gain H3K27ac in PND8-Ki67^+^, PLX-Ki67^+^ and CPZ-Ki67^+^ microglia at statistically significantly levels compared to Ki67^-^ microglia from healthy adult mice. **(H)** Average H3K27ac abundance over the - 750 to 750 bp region centered on promoter-distal distal elements that gain or lose acetylation in Ki67^+^ microglial subsets and Ki67^-^ microglia from healthy adult mice. **(I)** Venn diagram depicting overlap of among promoter-distal regions that gain or lose H3K27ac in PND8-Ki67^+^, PLX-Ki67^+^ and CPZ-Ki67^+^ microglia at statistically significantly levels compared to Ki67^-^ microglia from healthy adult mice.

Comparison of Ki67^-^ and Ki67^+^ microglia isolated from CPZ mice suggested that indeed the latter subset is engaged in proliferating activity. Indeed, among the 439 regions that are more acetylated at H3K27 in CPZ-Ki67^+^ microglia, there was a significant enrichment for regions the encompass the TSS of genes associated with cell division (Fig. 5C, D). Using principal component analysis (PCA), we next compared the genome-wide profile of H3K27ac of the CPZ microglial subsets with those of Ki67^-^ Adult-Basal microglia and Ki67^+^ microglia from PND8 neonates and PLX mice (Fig. 5E). Notably, CPZ-Ki67^+^ replicates clustered between CPZ-Ki67^-^ and PLX-Ki67^+^ microglia. This is coherent with both the brain microenvironment providing similar important regulatory input to both Ki67^-^ and Ki67^+^ microglia in the CPZ model, and the observation that PLX-Ki67^+^ microglia, but not PND8-Ki67^+^ microglia, possess an enhanced immune/inflammatory phenotype (Fig. 2B, C)^28^. Note that the very low prevalence of PLX-Ki67^-^ microglia did not allow for generation of ChIP-seq libraries of quality consistent with that of the other subsets and were thus not included.

We next investigated abundance of H3K27ac at promoter regions of C1 and C2 genes and promoter-distal regulatory elements to gain insights into the state of activity of these regions in CPZ-Ki67^+^ microglia. Overall, the H3K27ac signal correlated better amongst the three Ki67^+^ subsets than any of these with Ki67^-^ microglia from the Adult-Basal condition (Fig. 5F, S5A). The H3K27ac signal correlation in Ki67^-^ microglia from the healthy adult brain was also relatively more dissimilar at C2 gene promoters than at C1 promoters (Fig. S5A). Furthermore, a higher proportion of C2 gene promoters consistently gained H3K27ac across all three Ki67^+^ subsets than C1 promoters or a subset of randomly selected promoters (Fig. 5G). Finally, H3K27ac abundance at promoter-distal regulatory elements was less correlated among the Ki67^+^ microglia subsets at *Common-Up* regions defined earlier than at C1 or C2 promoters (Fig. 5H, I; S5B). This latter result is also concordant with promoter-distal elements activity being subject to important context-dependent signaling cues.

Collectively, these results confirm that a strong gain of activity, as measured by increased local deposition of the H3K27ac mark, occurs at C2 gene promoters during microglial proliferation. Notably, they also extend this mode of regulation associated with microglial proliferation to a context of inflammatory response associated with demyelinating brain lesions. Finally, they corroborate that microglial proliferation may proceed in absence of extensive coordinated gain of activity of large ensembles of promoter-distal regulatory elements.

## Discussion

The present study provides evidence for a transcriptional program whose proper regulation and activation may be essential to enable efficient microglial proliferation. Furthermore, this program integrates itself with the various other programs of gene regulation active in a given setting, which are subject to the regulatory influence of microglia’s surrounding brain microenvironment. Thus, our data does not support the hypothesis that microglia must attain a unique signature of gene expression or epigenomic landscape in order to proliferate. Given the alternative, the integration of the cell cycling gene program with additional, context-dependent transcriptional programs may help maintain their cell functions as they enter and proceed through the cell cycle process, or promptly re-engage them upon cell cycle exit. Ultimately, this could help the microglial population coordinate and stabilize its activities in promoting brain development, functions, repair and fight against infections.

At least two different modes of regulation possibly coordinate expression levels of genes required for microglial proliferation. Of interest, these two modes are also each linked to the regulation of distinct subsets of genes associated with dedicated cellular processes. A first mode augments the transcription of a group of genes that already expressed in non-proliferating G0 microglia and that are implicated in translation and ribonucleoside triphosphate metabolism (C1 genes). While these processes are not specific to cell cycling, they are nonetheless essential to its efficient regulation. Indeed, protein translation is a critical checkpoint of protein abundance dynamically regulated during cell cycle and cells must enhance synthesis of nucleotides prior to DNA synthesis^31,32^. In contrast, the second mode promotes the de novo expression of genes linked to various phases of cell cycling, including DNA replication and mitotic cell process (C2 genes). These include *Top2a*, *Lig1*, *Mki67*, *Cdc20*, among others. We note that the very strong induction of expression of C2 genes highlights how critical transcription as a checkpoint for regulation of microglial proliferation is.

These distinct modes of transcriptional control may operate through a two-tier mechanism involving combinatory activity of pan-regulators and mode-specific regulators. Based on our results, Klf/Sp, Nfy and Ets factors potentially act in broad capacity towards the transcriptional regulation a large array of C1 and C2 genes requires for microglial proliferation. Indeed, both Klfs/Sp1, Nfy and Ets motifs are highly prevalent at both C1 and C2 gene promoters and at promoter-distal regulatory elements that gain H3K27ac during microglial cell cycling. How the different family members of these genes modulate these different regulatory regions remains to be investigated. We note that activators Klf2, 4, 6 and 7, Sp1, NfyA-C, multiple Ets family members, and repressors Klf3 and 12, and activators/repressors Sp3-4^33–35^, are all reliably expressed in microglia (Fig. 3H). This observation not only suggests complex parallel regulation, but it may also provide a substantial level of redundancy with respect to the contribution of these factors. For example, genetic deletion of *Klf4* does not disrupt microglial cell proliferation during post-natal brain development and/or population in mice^36^. Such feature underlies how fundamental the effective input of these factors may be to ensure proper gene expression required for microglial cell cycling.

Transcriptional regulation to ensure efficient microglial proliferation requires additional input however. To this end, promoters of C1 but not C2 genes are enriched for motifs recognized by transcription factors Srebf2/E-box, Atf family members, and Yy1. Furthermore, the Nrf1 motif is frequently detected at C1 genes in both PND-Ki67^+^ and PLX-Ki67^+^ subsets. As Nrf1 is a major transcriptional regulator of genes implicated in mitochondrial biogenesis and functions^37,38^, this observation is coherent with the enhanced transcription of genes pertaining to these processes found within the C1 cluster. Alternatively, C2 gene promoters may require additional input from E2f and Lin54 factors for transcriptional activation. These have also been extensively characterized as fundamental regulators of gene expression necessary for cell cycling^39,40^. Furthermore, the strong upregulation of activators E2f1 and E2f2 suggests that transcriptional induction of these factors may be a critical checkpoint of the microglial cell cycling process (Fig. 3H). Once sufficiently expressed, these factors may in turn cooperate with Lin54, Nfy, Ets factors, and pro-transcription Klfs at promoters of C2 genes to transcribe the full ensemble of genes required for microglial proliferation. We note that such promoter-centric mechanism may also help explain the relatively large proportion of C2 promoters that gain H3K27ac in proliferating microglia across the conditions sampled here (e.g., Fig. 5G).

Additional evidence for a promoter-centric mechanism of gene regulation for microglial proliferation comes from the analysis of H3K27ac abundance at promoter-distal regulatory elements. Indeed, the overall number of elements that gained H3K27ac as a function of the proliferative process was relatively low, representing only 5.6% and 10.5% of the total number of promoter-distal elements for each of PND8-Ki67^+^ and PLX-Ki67^+^ subsets, respectively (Fig. 4A, B). This suggests that a low fraction of promoter-distal elements become increasingly more active during proliferation. Moreover, only Pu.1 and Klf were represented in over 15% of the elements that gained H3K27ac in both subsets. Overall, this may be indicative of a relatively minor contribution of promoter-distal elements in regulating cell cycling gene expression. That being said, it is possible that heterogenous, context-dependent signaling cues activate distinct promoter-distal regulatory elements, which in turn provide through chromatin folding regulatory input that converge on the same set of target genes. Future work leveraging chromatin conformation assays may help resolve this point.

Exactly how the transcription factors implicated in this study are regulated and the type of regulatory input that they provide to the RNA polymerase II transcriptional complex remain to be determined. Evidence indicates a fundamental role for signaling pathways and enzymatic complexes downstream of Csf1r^12,41^. However, as suggested by our data, other signaling pathways must provide critical input as well to properly adjust expression of key genes with microglial cells’ flexible needs, which are inherently context-dependent. Perhaps a strong illustration such *needs* is the observation that multiple transcripts coding for proteins required for cholesterol and fatty acid biosynthesis, including *Dhcr24* and *Sqle* mRNAs, are significantly more expressed in PLX-Ki67^+^ microglia than in PND8-Ki67^+^ microglia (Fig. 2B). This suggests that cholesterol sensing mechanisms provide important quantitative regulatory input onto mechanisms of gene expression. In addition, the extent to which context-specific signals provide effective, modifying unique input is not well understood. For example, whether the potentially strong AP-1 and Pu.1-Irf8 ternary complex activity present in Ki67^+^ microglia in the PLX model (Fig. S4B), but not in PND8 proliferating microglia, modulate the transcription of cell cycling genes remains to be determined.

The extent to which the transcriptional mechanisms unraveled here are similarly involved in mediating cell cycling required to sustain the homeostatic microglial population in the adult brain remains to be determined. At the population level, estimates put at < 2.5% the proportion of microglia that are engaged in a state of proliferation in the healthy adult mouse brain^4,5^. Such small percentage severely limits our ability to efficiently study these cells with the methodology used here. However, the robust deposition of H3K4me3 at promoters of C1 and C2 genes detected in Ki67^-^ microglia isolated from the healthy adult brain suggests that these genes are primed for transcriptional activation^26^. This observation provides tentative evidence that these promoters could be highly permissive for efficient transcription in most cases of microglial proliferation, including perhaps under homeostatic condition.

In sum, this study provides one of the first comprehensive characterization of the epigenomic/transcriptional mechanisms underlying microglial cell cycling. Given this, multiple questions arise from this work. Among them, we note that glial senescence, which is associated with a cell’s compromised ability to divide, has been linked to the pathophysiology of neurodegenerative diseases like Alzheimer’s disease^42^. Thus, it would be of high interest to determine how such defects may implicate the epigenomic and transcriptional mechanisms identified in this work. The degree to which our observations can be extended to human microglia will need to be resolved as well. Finally, we note that the experimental models and approaches that we established here provide a robust methodology to investigate at great resolution fundamental principles of cell proliferation in terminally differentiated cells in vivo. As such, they are well-suited for future work that will expand on our understanding of intricate signaling, epigenetic, and transcriptional mechanisms underlying cell cycling in complex physiological settings.

## Supporting information

Table S1. RNA-seq data associated with Figure 1.

Table S2. RNA-seq data associated with Figure 2.

Table S3. RNA-seq data associated with Figure 3.

## General

We thank Sophie Vachon for assistance with mouse colony management, and Nataly Laflamme for assistance with the cuprizone model of brain demyelination. We also thanks Dr. Johannes Schlachetzki for valuable comments on the manuscript. PLX5622 was provided under Materials Transfer Agreement by Plexxikon Inc.

## Author contributions

SB and DG conceived the study. SB ad AMX performed mice experiments and analyzed data. FDG analyzed data. SF performed experiments and analyzed flow cytometry data. SR contributed financially to this study and participated to data interpretation and manuscript preparation. DG generated sequencing libraries, analyzed data, and wrote the manuscript with assistance from SB, AMX, FDG and SR. DG also supervised all aspects of the project.

## Material and Methods

### Mice

C57BL/6J (Jackson no.000664) and Mki67^tm1.1Cle^ (Ki67-RFP; Jackson no.029802) were originally purchased from Jackson Laboratory and used to establish colonies at the animal research facility of Centre de Recherche du Centre Hospitalier Universitaire de Québec-Université Laval. For this study, only male mice were used. All adult mice were of 2 to 3.5 months of age at sacrifice. Mice had ab libitum access to food and water. All animal procedures were approved by Université Laval’s animal care and ethics committee (CPAUL3) and performed in compliance with ethical regulations and guidelines of the Canadian Council on Animal Care.

### PLX5622 treatment

To deplete microglia in the adult mouse brain, mice were fed PLX5622 (Plexxikon) formulated into AIN-76A chow pellets from Research Diets Inc (1200 PPM). Mice were given ab libitum access to PLX5622 chow diet for 17 consecutive days.

### Cuprizone diet

Cuprizone (Sigma-Aldrich) was mixed with regular ground chow (0.2% wt/wt) and fed ab libitum to mice for 4 weeks^30^. Diet was changed every 2 days.

### Microglia isolation

Mice were first euthanized with an overdose of ketamine-xylazine administered by intra-peritoneal injection and subsequently perfused transcardially with PBS. Brain were then removed from the skull and immediately homogenized in a buffer solution (HBSS (Life Technologies, 14175-095), 1% BSA (Fisher), 1mM EDTA (Invitrogen), 1 mM sodium butyrate (Sigma B5887-1G), 1 μM Flavoperidol (Sigma F3055) by gentle mechanical dissociation on ice, using a 2 ml polytetrafluoroethylene pestle (Wheaton, 358026) as performed in Gosselin et al. 2014^17^. Microglia were then enriched by Percoll (GE) gradient, washed and labeled with Live/Dead yellow dead stain (ThermoFisher), then incubated 1:100 with a CD16/CD32 receptor blocking antibody (1:50, eBioscience 14-0161, clone 93) for 20 min, and then incubated 1:100 with antibodies directed against cell surface proteins CD11b-BV421 (BioLegend 101251, clone M1/70), CD45-APC (BioLegend 103112, clone 30-F11), and CD44-APC-Cy7 (BioLegend 103028, clone IM7). For all ChIP-seq experiments, microglia were then further fixed with 1% PFA for 10 min at RT, permeabilized with reagents from ebioscience Foxp3 Transcription Factor staining buffer set (ThermoFisher 00-5523-00) and stained with anti-Ki67-Alexa 488 antibody (BD Pharmingen 561165, clone B56). Microglia were either analyzed with BD LSR or sorted with BD Aria II instruments (100 μm nozzle). Flow cytometry results were analyzed using FlowJo software (Tree Star).

### RNA-seq library preparation: Isolation and fragmentation of poly(A)

For RNA-seq, we generated 2 independent biological replicates per conditions (n=2). Following sorting, 200,000 microglia were pelleted and put into 100 μl lysis/Oligo d(T) Magnetic Beads binding buffer (100 mM Tris-HCl ph 7.5, 500 mM LiCl, 10 mM EDTA pH 8.0, 1% LiDS, 5 mM DTT, in water) and stored at −80°C until processing. For human brain cortical RNA, 500 ng of RNA was diluted with proper volume of 2x lysis/ Oligo d(T) Magnetic Beads binding buffer to a final concentration of 1x lysis/Oligo d(T) Magnetic Beads binding buffer. For sequencing library preparation, samples were incubated with 20 μl Oligo d(T) Magnetic Beads (NEB, S1419S) in PCR strips for selection of polyadenylated (poly(A)) mRNA as follows: 2 min at 65 °C on a PCR cycler and then 10 min at room temperature (RT). Samples were then placed on a collection magnet at RT, and washed serially with 180 μl of washing buffer 1 (RNA-WB1; 10 mM Tris-HCl pH 7.5, 0.15 M LiCl, 1 mM EDTA pH 8.0, 0.1% LiDS, 0.1% Triton X-100, in water) and then washing buffer 3 (RNA-WB3; 10 mM Tris-HCl, 0.15 M NaCl, 1 mM EDTA pH 8.0, in water). RNA was eluted from Oligo (t) beads by resuspending the beads in 50 μl of elution buffer (EB; 10 mM tris-HCl pH 7.5, 1 mM EDTA pH 8.0, in water) followed by incubation at 80°C for 2 min on a PCR cycler. The samples were then placed back on the magnet, and the elution buffer supernatant containing the poly(A) RNA was carefully collected and placed on ice.

A second round of poly(A) RNA selection was then performed. Eluted Oligo d(T) beads were first washed on a magnet with 150 μl EB and then 150 μl 2X Oligo d(T) binding buffer (2x DTBB; 20 mM Tris-HCl pH 7.5, 1 M LiCl, 2 mM EDTA pH 8.0, 1% LiDS, 0.1% Triton X-100) and resuspended in 50 μl of 2x DTBB. Resuspended Oligo d(T) beads were then added to isolated RNA and incubated at 65 °C and at RT as described above. Beads were then placed on a magnet and washed once with 150 μl of RNA-WB1 and then with 30 ul of 1x First-Strand buffer (made from 5x stock; Life Technologies) and transferred in a new PCR strip. Beads were collected on a magnet and resuspended in 10 ul of 2x SuperScript III buffer (Life Technologies, 1x First-Strand buffer, 10 mM DTT in water). Resuspended beads were then incubated at 94 °C for 9 min to fragment poly(A) RNA. Beads were then placed back on a magnet, and the supernatant containing poly(A) RNA was carefully collected and placed on ice.

### RNA-seq library preparation: Synthesis of cDNA

RNA in 2X SuperScript III buffer was incubated for 1 min at 50 °C on a PCR cycler with 2.5 μl of the following mix: 1.5 μg Random Primer (Life Technologies 48190-011), 10 μM Oligo d(T) (Life Technologies 18418020), 10 units SUPERase-In (Ambion AM2696), 4 mM dNTP mix (Life Technologies 18427088), in water. Samples were then immediately placed on ice for 5 min. First-strand synthesis was then performed by incubation at 25 °C for 10 min and 50 °C for 50 min on a PCR cycler with 7.6 μl of the following mix: 0.2 μg actinomycin (Sigma A1410), 13.15 mM DTT (Life Technologies), 0.026% Tween-20, 100 units SuperScript III (Life Technologies kit 18080-044), in water. After incubation, RNA/DNA complexes were isolated by adding 36 μl of Agencourt RNAClean XP beads (Beckman Coulter A63987) and incubated for 10 min at RT and then 10 min on ice. Samples were then placed on a magnet and beads were washed twice with 150 μl of 75% EtOH. Following washings, beads were air-dried for 10-12 min and eluted with 10 μl of water.

Second-strand synthesis was then performed. RNA/DNA samples in 10 μl of water were incubated for 2.5 hours at 16 °C with 5 μl of the following mix: 3x Blue Buffer (Enzymatics), 1.0 μl PCR mix (Affymetrix 77330), 2.0 mM dUTP (Affymetrix 77206), 1 unit RNAseH (Ezymatics Y9220L), 10 units DNA Polymerase I (Enzymatics P7050L), 0.03% Tween-20, in water). DNA was then purified by addition of 1.5 μl Sera-Mag SpeedBeads (Thermo Scientific, 651520505025), resuspended in 30 μl 20% PEG 8000/2.5 M NaCl, incubated at RT for 15 min and placed on a magnet for two rounds of bead washing with 80% EtOH. Beads were then air dried for 10-12 min and DNA was eluted from the beads by adding 40 μl of water. Supernatant was then collected on a magnet and placed on ice or stored at −20°C until DNA blunting, poly(A)-tailing, and adapter ligation (see below).

### Chromatin immunoprecipitation

Chromatin immunoprecipitation for histone modification H3K4me3 and H3K27ac and was performed as follows, using ~500,000 microglia per assay and n=2 independent biological replicates per conditions. First microglia were briefly thawed on ice and lysed by incubation in 1 ml of lysis buffer (0.5% IGEPAL CA-630, 10 mM HEPES pH 7.9, 85 mM KCl, 1 mM EDTA pH 8.0, in water) for 10 min on ice. Lysates were centrifuged at 800 RCF for 5 min at 4 °C, and pellets were resuspended in 200 μl of sonication/ immunoprecipitation buffer (10 mM Tris-HCl pH 7.5, 100 mM NaCl, 0.5 mM EGTA, 0.1% Deoxcycholate, 0.5% Sarkosyl, in water). Sonication was performed with a Bioruptor Standard Sonicator (Diagenode) and consisted of two rounds of 15 min each, alternating stages of 30 sec “sonication-on” with 60 sec “sonication-off”. Twenty-two microliters of 10% Triton-X were then added to samples (1% final concentration) on ice, and lysates were cleared by centrifugation for 5 min at 18,000 RCF at 4 °C. Two microliters of supernatant were then set aside for input library sequencing controls. Supernatants were then immunoprecipitated on a rotator for two hours at 4 °C with antibody pre-bound to 17 μl Protein A Dynabeads (Life Technologies 10001D). The following antibodies were used: H3K4me3 (Millipore, 04-745), 2.0 μl from stock per sample; H3K27ac (Active Motif, 39685), 2.5 μg per sample.

Immunoprecipitates were washed three times each on ice with ice-cold wash buffer I (150 mM NaCl, 1% Triton X-100, 0.1% SDS, 2 mM EDTA pH 8.0 in water), wash buffer III (10 mM Tris-HCl, 250 mM LiCl, 1% IGEPAL CA-630, 0.7% Deoxycholate, 1mM EDTA in water) and TET (10 mM Tris-HCl pH 7.5, 1 mM EDTA pH 8.0, 0.1% Tween-20, in water) and eluted with 1% SDS/TE at RT in a final volume of 100 μl.

Reverse-crosslinking was then performed on immunoprecipitates and saved input aliquots. First, 6.38 μl of 5 M NaCl (final concentration 300 mM) was added to each sample immersed in 100 μl SDS/TE. Crosslinking was then reversed by overnight incubation at 65 °C in a hot air oven. Potentially contaminated RNA was then digested for one hour at 37 °C with 0.33 mg/ml RNase A, proteins were digested for one hour at 55 °C with 0.5 mg/ml proteinase K, and DNA was extracted using Sera-Mag SpeedBeads (Thermo Scientific, 6515205050250).

### RNA-seq and ChIP-seq final library preparation

Sequencing libraries were prepared from recovered DNA (ChIP) or generated cDNA (RNA) by blunting, A-tailing, and adapter ligation as previously described using barcoded adapters (NextFlex, Bioo Scientific)^17,43^. Prior to final PCR amplification, RNA-seq libraries were digested by 30 min of incubation at 37°C with Uracil DNA Glycosylase (final concentration of 0.134 units per μl of library volume; UDG, Enzymatics G5010L) to generate strand-specific libraries. Libraries were PCR-amplified for 12-15 cycles and size selected for fragments (200-400 bp for ChIP-seq, 200-500 for RNA-seq) by gel extraction (10% TBE gels, Life Technologies EC62752BOX). RNA-seq and ChIP-seq librairies were single-end sequenced for 76 cycles on an Illumina HiSeq 4000 (Illumina, San Diego, CA) according to manufacturer’s instruction.

### Assay for Transposase-Accessible Chromatin-sequencing (ATAC-seq)

100,000 isolated microglia were lysed in 50 μl lysis buffer (10 mM Tris-HCl ph 7.5, 10 mM NaCl, 3 mM MgCl_2_, 0.1% IGEPAL, CA-630, in water) on ice and nuclei were pelleted by centrifugation at 500 RCF for 10 min. Nuclei were then resuspended in 50 μl transposase reaction mix (1x Tagment DNA buffer (Illumina 15027866), 2.5 μl Tagment DNA enzyme I (Illumina 15027865), in water) and incubated at 37°C for 30 min on a PCR cycler. DNA was then purified with Zymo ChIP DNA concentrator columns (Zymo Research D5205) and eluted with 10 μl of elution buffer. DNA was then amplified with PCR mix (1.25 μM Nextera primer 1, 1.25 μM Nextera primer 2-bar code, 0.6x SYBR Green I (Life Technologies, S7563), 1x NEBNext High-Fidelity 2x PCR MasterMix, (NEBM0541)) for 9-11 cycles, run on an gel for size selection of fragments (160-290 bp), extracted from the gel and single-end sequenced for 76 cycles on a HiSeq 4000.

### Sequencing data analysis

#### Preprocessing

FASTQ files from sequencing experiments were mapped to the mouse mm10 reference genome. STAR (2.7.0f) with default parameters was used to map RNA-seq experiments^44^. Bowtie2 (2.3.4.1) with default parameters was used to map ChIP-seq and ATAC-seq experiments^45^. HOMER was used to convert aligned reads into “tag directories” for further analyses^46^.

#### RNA-seq

For differential expression analyses, a table read count including relevant samples was first created using the *analyzeRepeats.pl* program of HOMER (v4.11; http://homer.ucsd.edu/homer/) with the following parameters: *rna -noadj - condenseGenes -count exons -pc 3*. Non-coding RNA and pseudogenes were then removed to retain only protein-coding mRNAs and the resulting dataset was used as input file for HOMER’s *getDiffExpression.pl* program, which leveraged R/DESeq2 for to assess differential expression. For data interpretation, only genes with 16 or more normalized reads were considered. Alternatively, Transcript Per Million-normalized datasets were generated using the *analyzeRepeats.pl* program of HOMER with the following parameters: rna -TPM -condenseGenes -count exons -pc 3. The base-2 logarithm of the TPM values was taken after adding a pseudocount of 1 TPM to each gene.

#### H3K4me3, H3K27ac ChIP-seq regions calling and differential analyses

For each condition, the repertoire of genomic regions positive for presence of H3K4me3 or H3K27ac were first generated using HOMER’s *getDiffPeakReplicate.pl* program, using all replicates and related input of a condition of interest. Parameters specified for H3K4me3 were as follow: *-style histone*. For H3K27ac, parameters were: *-genome mm10 -region -size 500 -min Dist 1000 -L1*. For differential enrichment assessments, relevant regions files were then first merged together with HOMER’s *mergedPeaks.pl* program with *-size given* parameter, and tag counts of each replicate annotated to each region using HOMER’s *annotatePeak.pl* program with *mm10 -raw* parameters used. For H3K27ac, data matrix was then used as input for HOMER’s *getDiffExpression.pl* program, which leveraged R/DESeq2 to assess differential enrichment. For read count tables output, each experiment was normalized to 10 million total reads (*-simpleNorm* default parameter specified in *getDiffExpression.pl*). For data interpretation, only regions with 16 or more normalized reads were considered.

#### ATAC-seq accessible chromatin region identification

Regions of accessible chromatin were called using HOMER’s *findPeaks.pl* program with the *-style factor* parameter.

#### ChIP-seq histograms

Histograms for tag density were generated using HOMER’s *annotatePeaks.pl* program, using as input the merged regions .txt files of relevant samples generated with the *-size given* parameter specified. Region annotation for promoters was performed in *tss* mode using *-list <relevant NM list.txt>*, with *-size. −1000,500 hist 10 -norm 1e7* parameters specified. For promoter-distal regions, default mode was used with *-size −750,750 hist 10 -norm 1e7* parameters specified. Control promoter for H3K4me3 histograms (Fig. 3D-E) consisted of 881 promoters of genes randomly selected from a list of genes not modulated in PND8-Ki67^+^ or PLX-Ki67^+^ and that were matched for tag density to C2 genes TPM values obtained from Ki67^-^ Adult-Basal microglia. Control promoters for Fig. 6G consisted of 999 promoters of genes randomly selected from a list of promoters that display at least 16 normalized tags in Ki67^-^ Adult-Basal microglia and that were not part of C1 or C2 clusters.

#### Venn diagram – H3K27ac at genomic regulatory elements

H3K27ac regions of interest that gain H3K27ac in PND8-Ki67^+^, PLX-Ki67^+^ and CPZ-Ki67^+^ microglia at statistically significantly levels compared to Ki67^-^ microglia from healthy adult mice were recovered from *getDiffExpression.pl* output .txt file. Overlap was then assessed using R.

### De novo motif enrichment analyses

#### Promoters analyses

For promoter analyses, de novo motif enrichment analysis was performed on the area of open chromatin defined by ATAC-seq (i.e., ATAC-seq “peaks”) located within −1000 to +500 bp genomic regions of transcriptional start sites (TSS) of C1 and C2 genes. For this, C1 and C2 gene promoters annotated with genomic coordinates (see above) were intersected with ATAC-seq “peaks” using BEDOPS with --*ec* --*element-of-1* parameter specified. BEDOPS output file was used with HOMER’s *findMotifsGenome.pl* command with the following parameters: *-size 200 -mask*. Background sequences for differential enrichment assessment were generated randomly by the program.

#### Promoter-distal regulatory elements analyses

Promoter-distal elements were defined as H3K27ac-positive regions located outside of the −1000 to 500 bp regions surrounding a gene TSS. DNA motif enrichment analyses for promoter-distal regulatory elements was centered on ATAC-seq peaks located within elements of interest (see Results), identified using the BEDOPS and HOMER’s *findMotifsGenome.pl* programs combination described above.

### Gene ontology analyses

Metascape with default parameters (https://metascape.org/) was used to perform gene ontology analyses on groups of genes of interest (RefSeq identifiers)^47^. Terms associated with GO Biological processes were retained.

### Data visualization

The UCSC genome browser was used to visualize ChIP-seq and ATAC-seq data^48^.

### Data and material availability

Original sequencing FASTQ files from all sequencing experiments have been deposited on the Gene Expression Omnibus platform (GEO; series GSE166236).

