## Supplementary Figure S1 to S5 for "Context-dependent transcriptional regulation of microglial proliferation"

**This PDF file includes:**

Figures S1 to S5
Table S1 to S3 (Titles)

**Figure S1**

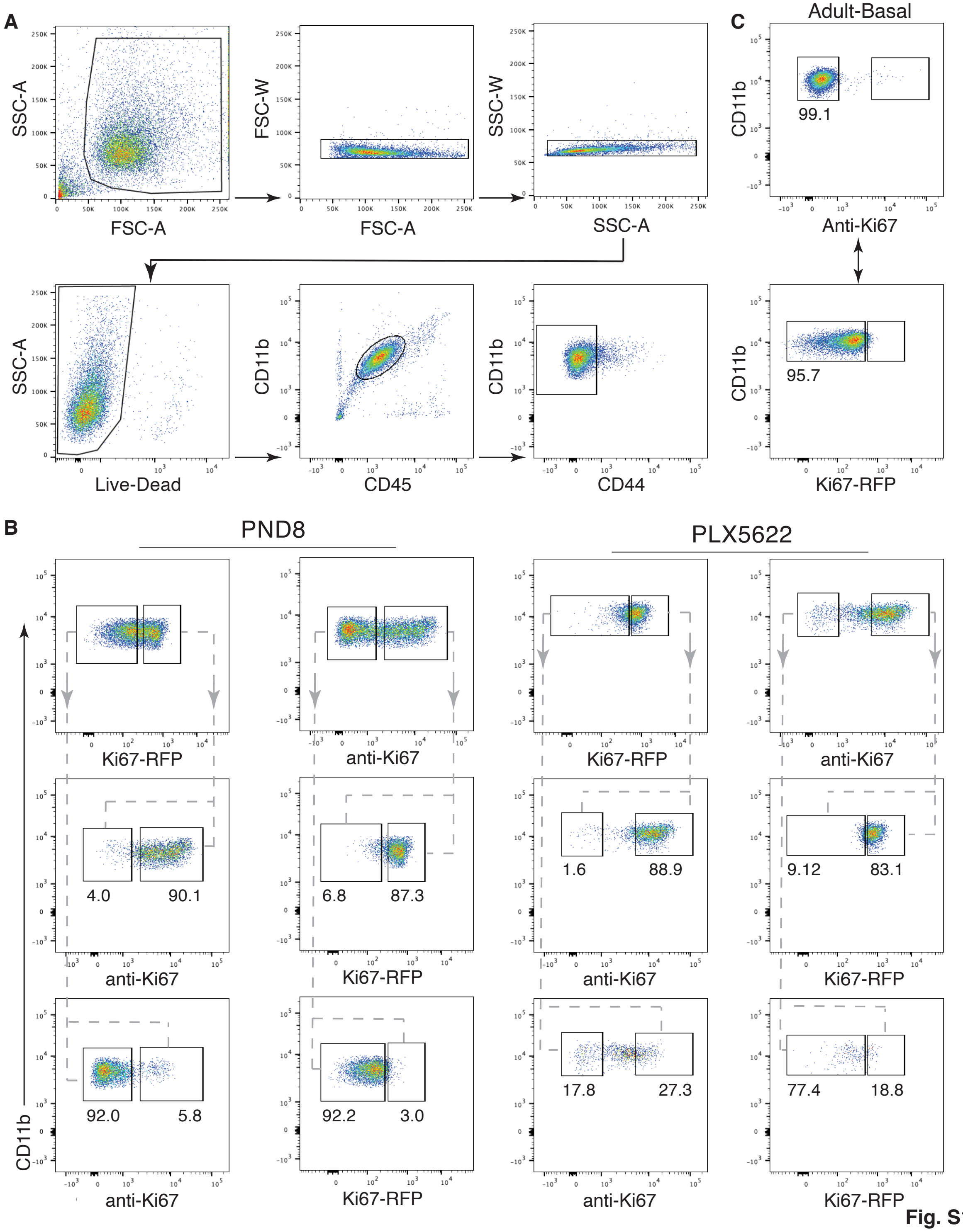


**Fig. S1**

**Figure S1. Gating strategy for isolation of Ki67$^{-}$ and Ki67$^{+}$ microglia. (A)** Microglia were defined as live-CD11b$^{+}$CD45$^{+}$CD44$^{Low}$ cells. Ki67 expression was then used to discriminate Ki67$^{+}$/proliferative from Ki67$^{-}$ microglia. **(B)** Concordance of Ki67 expression as revealed by RFP signal in Ki67-RFP mice and antibody staining using anti-Ki antibody. Numbers represent percentage of microglia in corresponding gate above, relative to parent gate. **(C)** Ki67 antibody staining of microglia isolated from adult Ki67-RFP mice.

### Figure S2

**C1 promoters**

(Motifs analysis centered on PLX-Ki67$^{+}$ ATAC-seq regions)

| Motif best match (score) | p-value | % target/ %bckg |
|---|---|---|
| Klf/Sp (0.91) | 1e-94 | 46/20 |
| Nfy (0.96) | 1e-73 | 23/7 |
| Ets (0.95) | 1e-59 | 25/10 |
| Nrf (0.96) | 1e-41 | 16/5 |
| Gfy (0.98) | 1e-27 | 5/0.8 |
| Atf1 (0.86) | 1e-25 | 17/8 |
| Yy1 (0.82) | 1e-24 | 18/9 |

**C2 promoters**

(Motifs analysis centered on PLX-Ki67$^{+}$ ATAC-seq regions)

| Motif best match (score) | p-value | % target/ %bckg |
|---|---|---|
| Nfy (0.95) | 1e-83 | 31/6 |
| Klf/Sp (0.92) | 1e-66 | 42/14 |
| Lin54 (0.91) | 1e-34 | 25/8 |
| E2f (0.86) | 1e-30 | 20/6 |
| Ets (0.94) | 1e-20 | 28/13 |
| Gfy (0.81) | 1e-18 | 5/0.5 |
| Nrf (0.89) | 1e-14 | 13/5 |

**Fig. S2**

**Figure S2. DNA Motifs analyses at C1 and C2 promoters.** De novo motifs enrichment was performed, centered on ATAC-seq-defined regions of accessible chromatin from PLX-Ki67$^+$ microglia included within the -1000 to + 500 bp regions surrounding TSS of C1 (left) and C2 (right) genes.

**Figure S3**

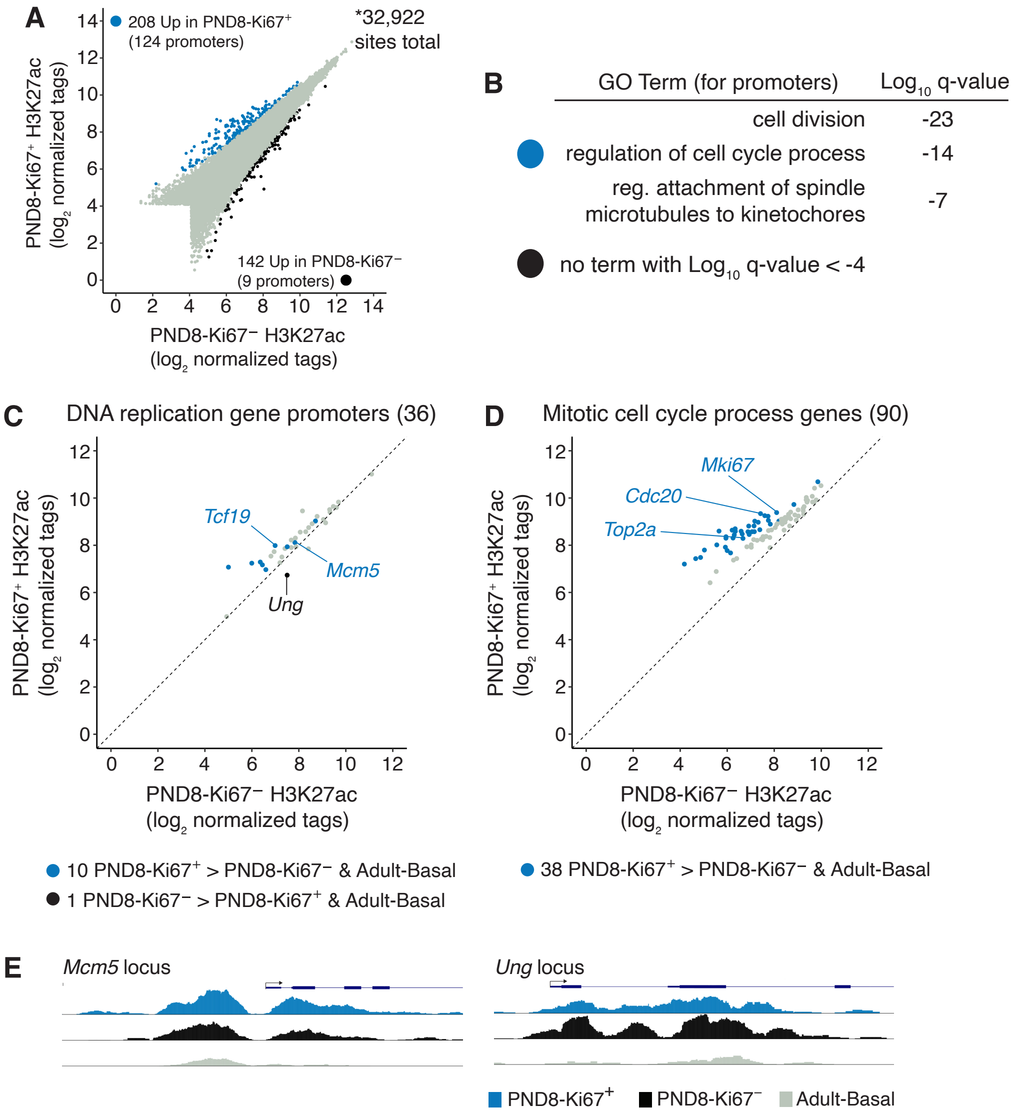


**Fig. S3**

**Figure S3. Genome-wide profiles of H3K27ac for PND8-Ki67$^-$ and PND8-Ki67$^+$ microglia. (A)** Comparison of genome-wide profile comparison of H3K27ac PND8-Ki67$^-$ and PND8-Ki67$^+$ microglia. **(B)** Gene ontology analyses for promoter regions significantly more enriched for H3K27ac in PND8-Ki67$^+$ vs PND8-Ki67$^-$ microglia, and vice versa. **(C)** H3K27ac abundance at promoters of DNA replication genes, comparing PND8-Ki67$^-$ and PND8-Ki67$^+$ microglia. Promoters with significantly higher H3K27ac signal are colored (see legend). **(D)** H3K27ac abundance at promoters of genes involved in mitosis, comparing PND8-Ki67$^-$ and PND8-Ki67$^+$ microglia. Promoters with significantly higher H3K27ac signal are colored (see legend). **(E)** UCSC genome browser visualization of H3K27ac signals for selected genes.

### Figure S4

**A**

**Common-Up** promoter-distal elements
(Motifs analysis centered on PLX-Ki67$^+$ ATAC-seq regions)

| Motif best match (score) | p-value | % target/ %bckg |
|---|---|---|
| Spi1 (0.93) | 1e-141 | 44/9 |
| JunB (AP-1; 0.98) | 1e-28 | 11/2 |
| Pu.1-Irf (0.93) | 1e-28 | 8/1 |
| Mef (0.90) | 1e-17 | 7/2 |
| Runx (0.96) | 1e-16 | 5/0.8 |
| Klf (0.87) | 1e-16 | 16/7 |
| Ctcfl (0.76) | 1e-14 | 5/1 |
| Smad3 (0.73) | 1e-12 | 5/1 |

**B**

**PND8-Ki67$^+$-biased Up** promoter-distal elements
(Motifs analysis centered on PND8-Ki67$^+$ ATAC-seq regions)

| Motif best match (score) | p-value | % target/ %bckg |
|---|---|---|
| Spi1 (0.94) | 1e-189 | 38/6 |
| Mef2 (0.97) | 1e-25 | 4/0.4 |
| Mafb (0.93) | 1e-23 | 12/4 |
| Runx (0.93) | 1e-19 | 12/5 |
| Ctcfl (0.89) | 1e-19 | 5/0.8 |
| Drmt1 (0.67) | 1e-16 | 8/2 |
| Elf3 (0.73) | 1e-16 | 7/2 |
| Plagl2 (0.76) | 1e-14 | 5/1 |

**PLX-Ki67$^+$-biased Up** promoter-distal elements
(Motifs analysis centered on PLX-Ki67$^+$ ATAC-seq regions)

| Motif best match (score) | p-value | % target/ %bckg |
|---|---|---|
| Spi1 (0.95) | 1e-534 | 56/14 |
| JunB (AP-1; 0.98) | 1e-162 | 19/4 |
| Pu.1-Irf (0.93) | 1e-97 | 12/3 |
| Mef2 (0.89) | 1e-37 | 4/0.9 |
| Atf (0.90) | 1e-35 | 10/4 |
| Runx (0.94) | 1e-30 | 27/18 |
| Ctcfl (0.91) | 1e-25 | 4/1 |
| Klf (0.95) | 1e-20 | 9/5 |

**C**

**PND8-Ki67$^+$-biased Down** promoter-distal elements
(Motifs analysis centered on Ki67$^-$-Adult-Basal ATAC-seq regions)

| Motif best match (score) | p-value | % target/ %bckg |
|---|---|---|
| Spi1 (0.96) | 1e-445 | 54/12 |
| Pu.1-Irf (0.85) | 1e-37 | 9/3 |
| Batf (AP-1; 0.98) | 1e-37 | 14/6 |
| Gata4 (0.76) | 1e-34 | 9/3 |
| Atf (0.91) | 1e-28 | 10/4 |
| Smad2 (0.89) | 1e-20 | 25/17 |
| Mef2 (0.84) | 1e-19 | 4/1 |
| Mxi1(Bhlh; 0.88) | 1e-18 | 20/12 |

**PLX-Ki67$^+$-biased Down** promoter-distal elements
(Motifs analysis centered on Ki67$^-$-Adult-Basal ATAC-seq regions)

| Motif best match (score) | p-value | % target/ %bckg |
|---|---|---|
| Spi1 (0.96) | 1e-654 | 54/11 |
| Maf (0.92) | 1e-54 | 22/11 |
| Tfec (0.94) | 1e-50 | 8/3 |
| Mef2 (0.96) | 1e-42 | 6/2 |
| Cebp (0.93) | 1e-26 | 4/1 |
| Ctcf (0.82) | 1e-20 | 2/0.4 |
| Runx (0.85) | 1e-18 | 4/1 |
| Atf (0.90) | 1e-18 | 7/4 |

**Fig. S4**

**Figure S4. DNA Motifs analyses at promoter-distal regulatory elements. (A)** Results for *Common-Up* regions, using ATAC-seq data from PLX-Ki67$^{+}$ microglia. **(B)** Results for PND8-Ki67$^{+}$-biased *Up* (left) and PLX-Ki67$^{+}$ -biased *Up* (right) promoter-distal elements. **(C)** Results for PND8-Ki67$^{+}$-biased *Down* (left) and PLX-Ki67$^{+}$-biased *Down* (right) promoter-distal elements.

### Figure S5

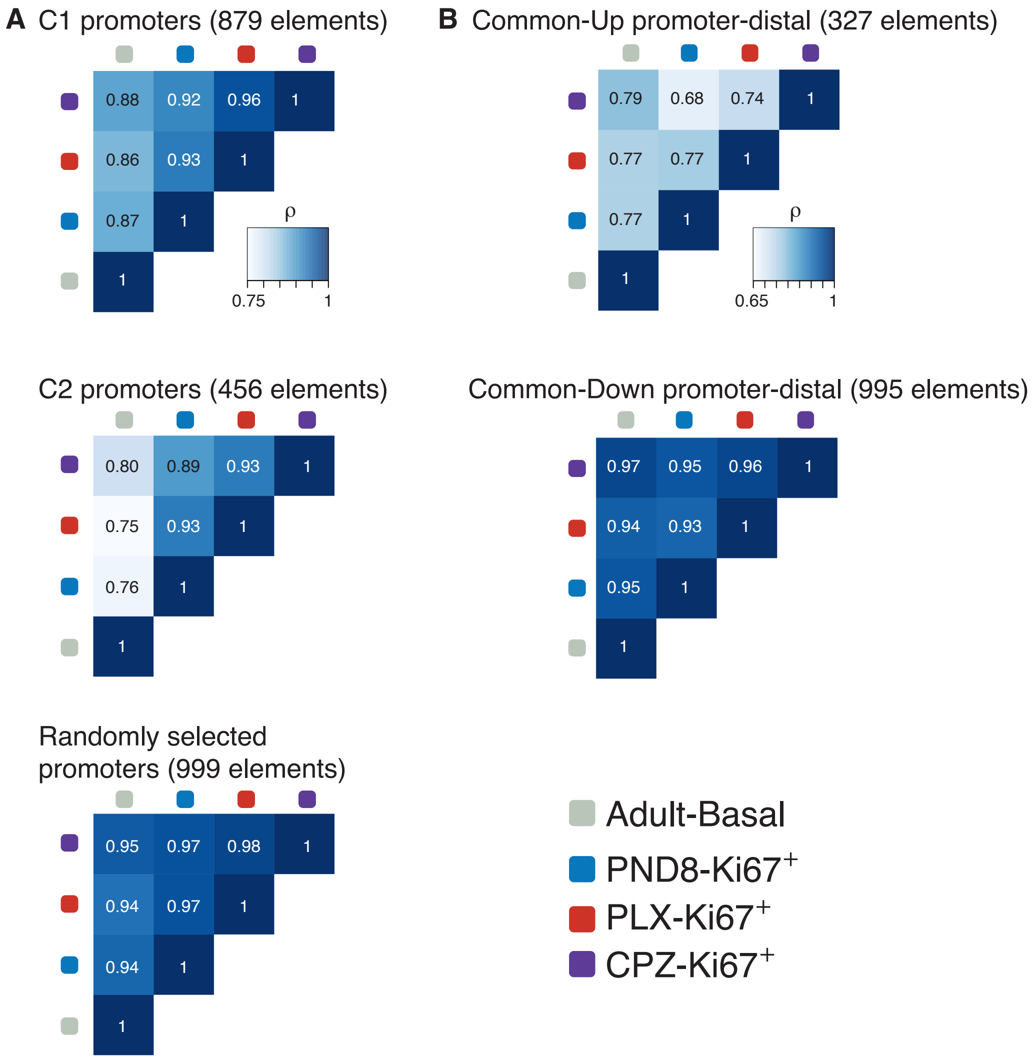


Fig. S5

**Figure S5. H3K27ac ChIP-seq signal correlation for various classes of regulatory elements.**

**(A)** Pearson correlation matrix of H3K27ac ChIP-seq signal over C1, C2, control gene promoters comparing different $Ki67^{+}$ microglial subsets and $Ki67^{-}$ microglia from healthy adult. **(B)** Correlations for *Common-Up* and *Common-Down* promoter-distal regulatory elements.

**Table S1. RNA-seq data associated with Figure 1.**

**Table S2. RNA-seq data associated with Figure 2.**

**Table S3. RNA-seq data for Cluster C1 and Cluster C2 genes, associated with Figure 3.**
